# Docking studies in targeting proteins involved in cardiovascular disorders using phytocompounds from *Terminalia arjuna*

**DOI:** 10.1101/2020.06.22.164129

**Authors:** Vikas Kumar, Nitin Sharma, Anuradha Sourirajan, Prem Kumar Khosla, Kamal Dev

## Abstract

*Terminalia arjuna* (Roxb.) Wight and Arnot (*T. arjuna*) commonly known as Arjuna has been known for its cardiotonic nature in heart failure, ischemic, cardiomyopathy, atherosclerosis, myocardium necrosis and also has been used in the treatment of different human disorders such as blood diseases, anaemia and viral diseases. Our focus has been on phytochemicals which do not exhibit any cytotoxicity and have significant cardioprotective activity. Since Protein-Ligand interactions play a key role in structure-based drug design, therefore with the help of molecular docking, we screened 19 phytochemicals present in *T. arjuna* and investigated their binding affinity against different cardiovascular target proteins. The three-dimensional (3D) structure of target cardiovascular proteins were retrieved from Protein Data Bank, and docked with 3D Pubchem structures of 19 phytochemicals using Autodock vina. Molecular docking and drug-likeness studies were made using ADMET properties while Lipinski’s rule of five was performed for the phytochemicals to evaluate their cardio protective activity. Among all selected phytocompounds, arjunic acid, arjungenin, and terminic acid were found to fulfill all ADMET rules, drug likeness, and are less toxic in nature. Our studies, therefore revealed that these three phytochemicals from *T. arjuna* can be used as promising candidates for developing broad spectrum drugs against cardiovascular diseases.

## Introduction

Cardiovascular diseases (CVD) are one of major cause of death in both developed and developing countries accounting approximately 31% of all deaths worldwide. Urbanization and changes in life style are main causes for increase of CVDs. It is expected that by the year 2030, more than 23.3 million people will die annually due to CVDs. It has been estimated that 17.7 million people died from CVDs in 2015, out of which, 7.4 million were due to coronary heart disease and 6.7 million were due to stroke. Cardiovascular diseases (CVDs) are a group of disorders of the heart and blood vessels including coronary heart disease, cerebrovascular disease, peripheral arterial disease, rheumatic heart disease, congenital heart disease, deep vein thrombosis and pulmonary embolism.

The renin-angiotensin system is an important regulatory system of cardiovascular and renal function. Any inhibition to this system, i.e. angiotensin converting enzyme inhibitors and angiotensin receptor has become a front-line treatments for hypertensive target organ damage and progressive renal disease [1]. High blood cholesterol level is one of the key factors for progression of Coronary artery disease (CAD) one of the key factors [2]. Inhibition to 3-hydroxy-3-methylglutaryl-coenzyme A reductase (HMGCR), a major cholesterol biosynthesis enzyme can result in decreasing the risk of coronary complications [3]. Braz *et al*. [4] reported that MAPK P38 plays a pivotal role in the signal transduction pathway mediating post-ischemic myocardial apoptosis and inhibition to MAPK P38 may attenuate reperfusion injury. Any stimulation of MAPK P38 induces intrinsic myocardial apoptosis activity and cellular signaling that results in reperfusion injury. β-Adrenergic receptors (β1AR, and β2AR) are G protein coupled receptors and are a promising tool that play regulatory function in the cardiovascular system (CVS) via signaling. It increases the Gα level, which activates βAR kinase (βARK) that affects and enhances the progression of heart failure (HF) through the activation of cardiomyocyte β-Adrenergic receptors [5]. C-reactive protein, among other systemic inflammatory mediators, has been widely accepted as a potent risk indicator, independently predicting future cardiovascular events. Evidence has cumulated to show that C-reactive protein levels are associated with different aspects of the cardiovascular risk spectrum [6]. An increased risk of cardiovascular events has been associated with the use of Non-steroidal anti-inflammatory drugs (NSAIDs), especially of COX-2 selective NSAIDs [7]. The synthetic drugs used for treatment of various cardiovascular diseases showed several side effects. In modern era, plants based medicinal system has become more popular because of low cost, easily availability, more effectiveness and lack of side effects.

*Terminalia arjuna* (Roxb) Wight and Arnot (*T. arjuna*) commonly known as ‘Arjuna’ has been used as cardiotonic in heart failure, ischemic, cardiomyopathy, atherosclerosis, myocardium necrosis [8-11] and also has been used in the treatment of different human disorders such as blood diseases, anaemia and viral diseases [12-15]. Several reports have studied the cardioprotective potential of *T. arjuna* stem bark using various models [16-21]; but mechanism of action of phytocompounds of *T. arjuna* against cardiovascular target proteins is still unclear. Mythili *et al*. [22] reported that arjunolic acid which is present in *T. arjuna* stem bark has protective effect against CsA-induced cardiotoxicity. To extend our research in cardiovascular disorders, we made a new approach to find the interaction between the phytochemical compounds from *T. arjuna* and synthetic drugs (Phosphocholine, Simvastatin, Rofecoxib, Carazolol, Lisinopril, Losartan, Losmapimod) and cardiovascular targets using molecular docking tools. Therefore, we investigated *in silico* docking analysis of cardiovascular disease targets with phytocompounds from *T. arjuna* tree.

## Methodology

### Ligand preparation

The 3D structures of selected phytocompounds and synthetic drugs were obtained from Pubchem in SDF format and converted to PDB format using the tool Open Babel [23]. The ligands were available in XML, SDF, JSON and ASN.1 format. Then the Gasteiger charges and rotatable bonds were assigned to the PDB ligands using Auto Dock Tool [24]. All rotatable bonds were allowed to move freely. The compound structures were energy minimized and considered for docking studies (Table 1).

**Table-1:** Major Phytocompounds present in various parts of *T. arjuna* for docking studies.

| Sr. No | Compounds/Drugs | Formula | Compound ID |
| --- | --- | --- | --- |
| 1 | Arjunetin | C<sub>36</sub>H<sub>58</sub>O<sub>10</sub> | 3052779 |
| 2 | Arjunic acid | C<sub>30</sub>H<sub>48</sub>O<sub>5</sub> | 15385516 |
| 3 | Arjunolic acid | C<sub>30</sub>H<sub>48</sub>O<sub>5</sub> | 73641 |
| 4 | Arjunone | C<sub>19</sub>H<sub>20</sub>O<sub>6</sub> | 14034821 |
| 5 | Arjungenin | C<sub>30</sub>H<sub>48</sub>O<sub>6</sub> | 12444386 |
| 6 | β-sitosterol | C<sub>29</sub>H<sub>50</sub>O | 222284 |
| 7 | Casuarinin | C<sub>41</sub>H<sub>28</sub>O<sub>26</sub> | 13834145 |
| 8 | Ellagic acid | C<sub>14</sub>H<sub>6</sub>O<sub>8</sub> | 5281855 |
| 9 | Ethylgallate | C<sub>9</sub>H<sub>10</sub>O<sub>5</sub> | 13250 |
| 10 | Gallic acid | C<sub>7</sub>H<sub>6</sub>O<sub>5</sub> | 370 |
| 11 | Luteolin | C<sub>15</sub>H<sub>10</sub>O<sub>6</sub> | 5280445 |
| 12 | Quercetin | C<sub>15</sub>H<sub>10</sub>O<sub>7</sub> | 5280343 |
| 13 | Terminic acid | C<sub>30</sub>H<sub>48</sub>O<sub>4</sub> | 355369274 |
| 14 | Catechin | C<sub>15</sub>H<sub>14</sub>O<sub>6</sub> | 73160 |
| 15 | Rutin | C<sub>27</sub>H<sub>30</sub>O<sub>16</sub> | 5280805 |
| 16 | Kaempferol | C<sub>15</sub>H<sub>10</sub>O<sub>6</sub> | 5280863 |
| 17 | Leucocyanidin | C<sub>15</sub>H<sub>14</sub>O<sub>7</sub> | 71629 |
| 18 | 3-O-Methylellagic acid 3'-rhamnoside | C<sub>21</sub>H<sub>18</sub>O<sub>12</sub> | 73157208 |
| 19 | Oleanolic acid | C<sub>30</sub>H<sub>48</sub>O<sub>3</sub> | 10494 |
| **Drugs** |  |  |  |
| 20 | Phosphocholine (PC) | C<sub>5</sub>H<sub>15</sub>NO<sub>4</sub>P<sup>+</sup> | 1014 |
| 21 | Simvastatin | C<sub>25</sub>H<sub>38</sub>O<sub>5</sub> | 54454 |
| 22 | Rofecoxib | C<sub>17</sub>H<sub>14</sub>O<sub>4</sub>S | 5090 |
| 23 | Carazolol | C<sub>18</sub>H<sub>22</sub>N<sub>2</sub>O<sub>2</sub> | 71739 |
| 24 | Lisinopril | C<sub>21</sub>H<sub>31</sub>N<sub>3</sub>O<sub>5</sub> | 5362119 |
| 25 | Losartan | C<sub>22</sub>H<sub>23</sub>ClN<sub>6</sub>O | 3961 |
| 26 | Losmapimod | C<sub>22</sub>H<sub>26</sub>FN<sub>3</sub>O<sub>2</sub> | 11552706 |

### Protein preparation

A total of seven proteins which were elated to several cardiovascular diseases were selected on the basis of literature survey (Table 2). The 3D crystal structures of selected target proteins were obtained from the RCSB protein data bank (http://www.pdb.org). All the proteins had co-crystallized ligands (X-ray ligand) in the binding site. The ligands enclosed in each protein structure were removed from the binding site and saved to a new file. Automated molecular docking was performed to find out molecular interaction and optimized geometry by using docking software AutoDock vina. All heteroatoms including water molecules, and definitions for symmetry were excluded from the file. Polar hydrogens were added to each protein and it was minimized by applying Kollman’s partial atomic charges. Minimized structure was saved in PDBQT file format that contains a protein structure with hydrogen in all polar residues. All docking parameters were set to default values on the basis of Lamarckian genetic algorithm principle [25].

**Table 2:**
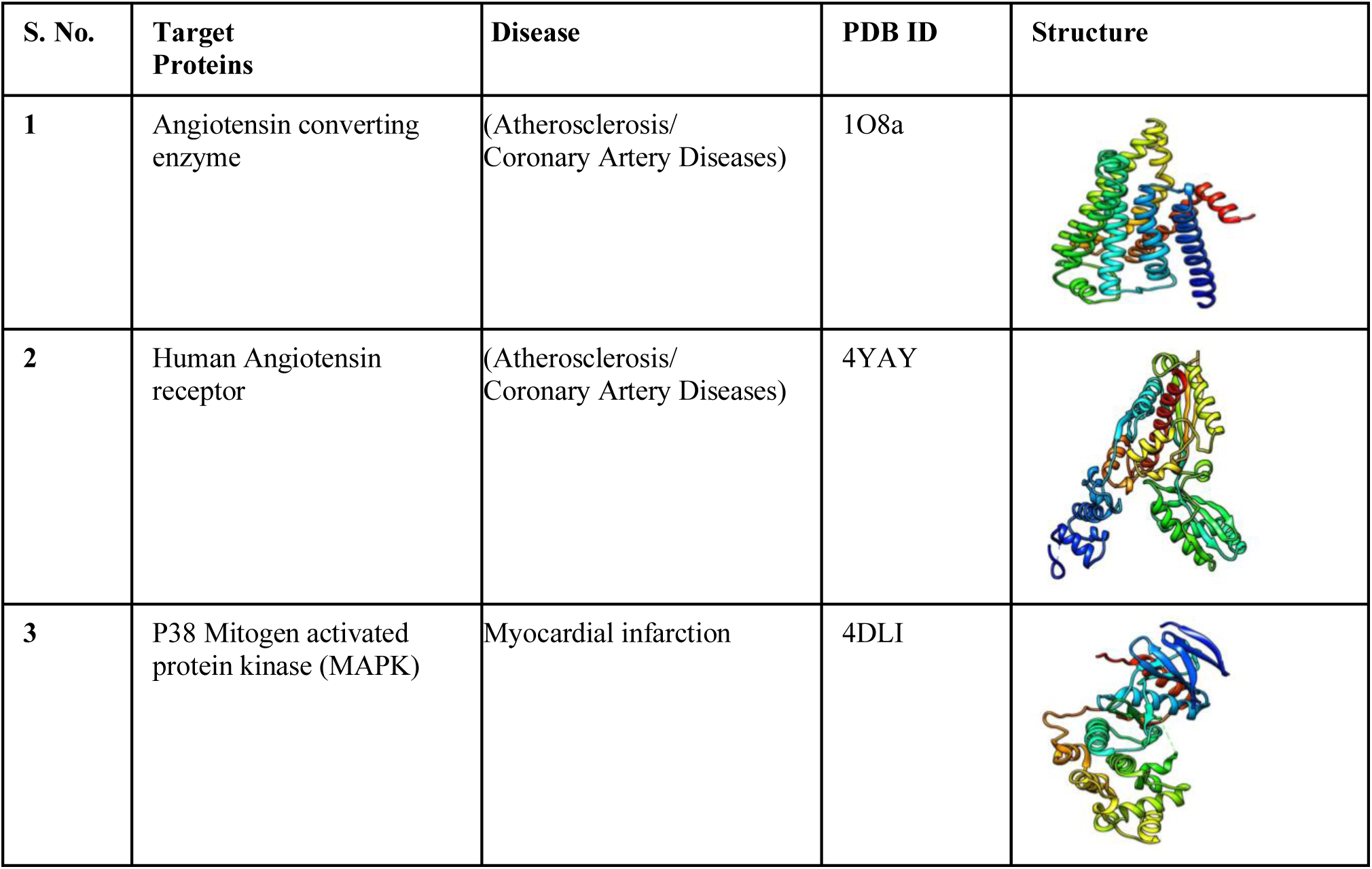
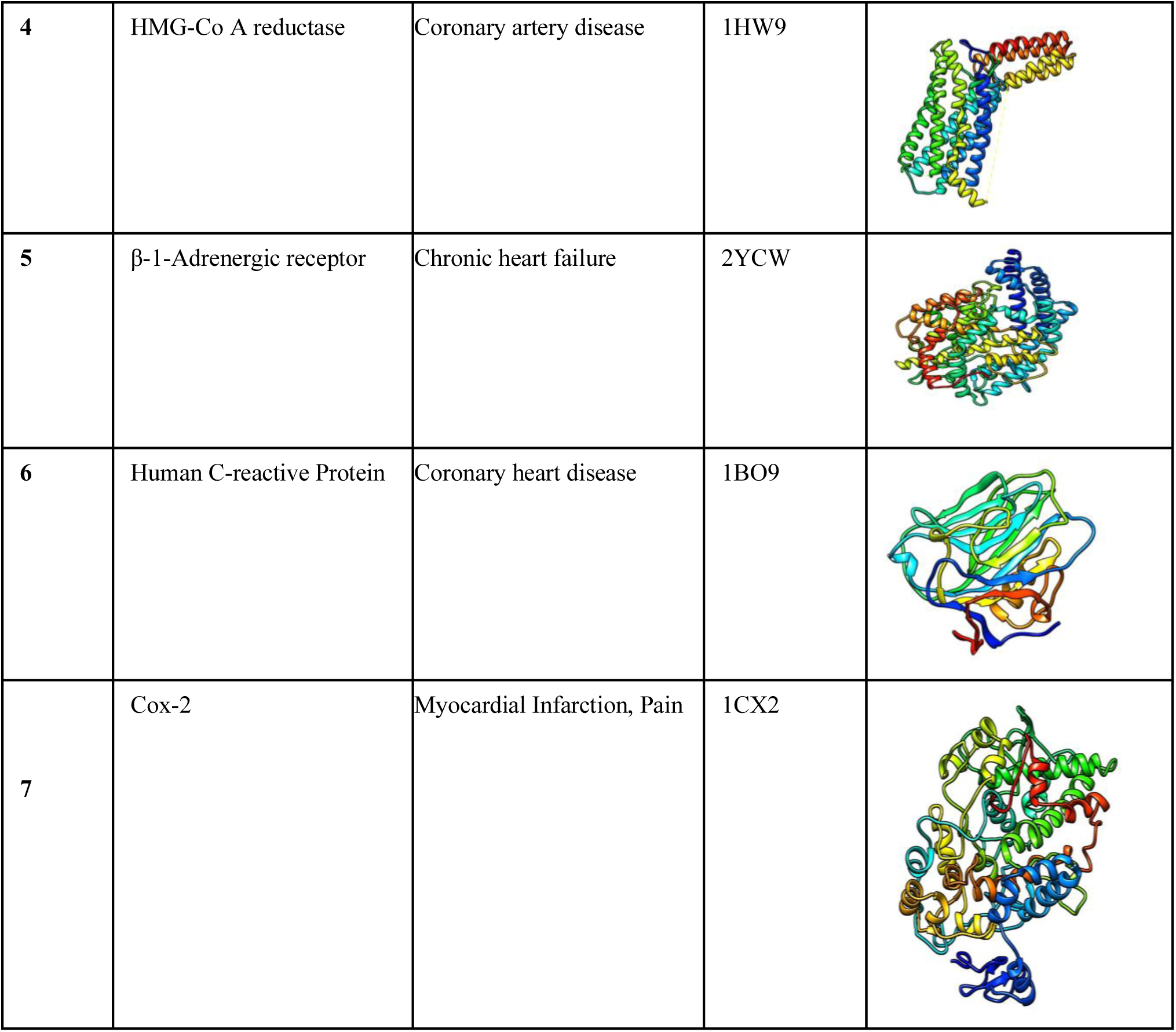
Targeted receptor proteins associated with cardiovascular disease along with structure.

| S. No. | Target Proteins | Disease | PDB ID | Structure |
| --- | --- | --- | --- | --- |
| 1 | Angiotensin converting enzyme | (Atherosclerosis/<br>Coronary Artery Diseases) | 1O8a |  |
| 2 | Human Angiotensin receptor | (Atherosclerosis/<br>Coronary Artery Diseases) | 4YAY |  |
| 3 | P38 Mitogen activated protein kinase (MAPK) | Myocardial infarction | 4DLI |  |
| 4 | HMG-Co A reductase | Coronary artery disease | 1HW9 |  |
| 5 | β-1-Adrenergic receptor | Chronic heart failure | 2YCW | |
| 6 | Human C-reactive Protein | Coronary heart disease | 1BO9 |  |
| 7 | Cox-2 | Myocardial Infarction, Pain | 1CX2 |  |

### Grid box generation

The grid parameter file of each protein was generated using AutoDock Tool. A grid-box was created in such a way that it covers the entire protein binding site and accommodates all ligands to move freely in it. The center of the active binding site was estimated from the structure and taken as the center of the grid-box.

### Molecular docking

The docking of selected ligands to the catalytic triad of protein was performed using AutoDock vina. Docking was performed to obtain a population of possible conformations and orientations for the ligand at the binding site. Polar hydrogen atoms were added to all the proteins and its non polar hydrogen atoms were merged by using the software. All bonds of ligands were set to be rotatable. All calculations for ligand flexible protein-fixed docking were performed using the Lamarckian Genetic Algorithm (LGA) method. The best conformation was chosen with the lowest docked energy, after the completion of docking search. Standard docking settings were used and the 10 energetically most favorable binding poses are outputted. The interactions between selected phytocompounds and target receptor were visualized by using UCHF chimera 1.8.1 [26] and LigPlot [27].

### ADMET and toxicity prediction

Absorption, distribution, metabolism, excretion, and toxicity (ADMET) screening of ligands helps to determine their absorption properties, toxicity, and drug-likeness. Ligand molecules saved in .smiles format and selected drugs (Phosphocholine, Simvastatin, Rofecoxib, Carazolol, Lisinopril, Losartan, Losmapimod)were uploaded on SWISSADME (Molecular Modeling Group of the SIB (Swiss Institute of Bioinformatics), Lausanne, Switzerland), admetSAR (Laboratory of Molecular Modeling and Design, Shanghai, China), and PROTOX webservers (Charite University of Medicine, Institute for Physiology, Structural Bioinformatics Group, Berlin, Germany) for ADMET screening. SWISSADME is a web tool used for the prediction of ADME and pharmacokinetic properties of a molecule. The predicted result consists of lipophilicity, water solubility, physicochemical properties, pharmacokinetics, drug-likeness, medicinal chemistry, and Brain or Intestinal Estimated permeation method (blood– brain barrier and PGP ± prediction) [28]. AdmetSAR provides ADMET profiles for query molecules and can predict about fifty ADMET properties. Toxicity classes are Category I containing compounds with LD_50_ values ≤50 mg/kg, Category II containing compounds with LD_50_ values >50 mg/kg; Category III (slightly toxic) containing compounds with 500 < LD50 ≤ 5000 mg/kg and safe chemicals (LD_50_ > 5000 mg/kg) are included in Category IV [29, 30]. PROTOX is a Rodent oral toxicity server predicting LD_50_ value and toxicity class of query molecule. The toxicity classes are as follows: (i) Class 1: fatal if swallowed (LD50 ≤5), (ii) Class 2: fatal if swallowed (55000) [31].

### Drug likeness calculations

Drugs scans were carried out to determine whether the phytochemicals fulfil the drug-likeness conditions. Lipinski’s filters using Molinspiration (http://www.molinspiration.com) were applied for examining drug likeness attributes as including quantity of hydrogen acceptors (should not be more than 10), quantity of hydrogen donors (should not be more than 5), molecular weight (mass should be more than 500 daltons) and partition coefficient log P (should not be less than 5). The smiles format of each of the phytochemical was uploaded for the analysis

## Results

### Molecular docking analysis

Docking results of selected phytocompounds with targeted proteins i.e. 1O8a, 4YAY, 4DLI, 1HW9, 2YCW, 1BO9 and 1CX2 showed that selected phytochemicals from *T. arjuna* had a good binding affinity and better binding modes than that of standard drugs against selected target receptors. All the selected 19 phytocompounds showed strong or comparative binding with all selected receptors. The data of binding energy of selected phytocompounds with target proteins is presented in table 3.

**Table 3:** Binding energy of 19 phytocompounds from *T. arjuna* against targeted protein receptors. Binding energy was expressed in terms of Kcal/mol.

| Compounds | 1B09 | 1HW9 | 1CX2 | 4YCW | 1O8a | 4YAY | 4dli |
| --- | --- | --- | --- | --- | --- | --- | --- |
| Arjunetin | -6.6 | -4.6 | -7.8 | -8.1 | -6.6 | -10.3 | -7.1 |
| Arjunic acid | -13.3 | -7.2 | -14.3 | -16.9 | -7.1 | -8.8 | -8.0 |
| Arjunolic acid | -12.7 | -7.4 | -14.2 | -16.8 | -5.2 | -10.1 | -7.4 |
| Arjunone | -8.1 | -4.2 | -7.0 | -9.5 | -5.1 | -8.7 | -7.1 |
| Arjungenin | -16.1 | -8.4 | -17.3 | -19.3 | -8.4 | -16.5 | -21.6 |
| β-sitosterol | -10.9 | -5.7 | -7.8 | -13.2 | -6.2 | -13.0 | -9.0 |
| Casuarinin | -15.4 | -12.0 | -19.4 | -24.0 | -14.7 | -21.0 | -17.0 |
| Ellagic acid | -7.9 | -4.6 | -9.3 | -10.6 | -6.7 | -10.2 | -11.8 |
| Ethyl gallate | -6.5 | -3.2 | -6.5 | -6.8 | -4.1 | -6.9 | -7.3 |
| Gallic acid | -6.3 | -2.7 | -6.0 | -6.7 | -4.2 | -6.3 | -6.8 |
| **Luteolin** | -9.3 | -4.6 | -7.8 | -10.3 | -5.9 | -9.0 | -8.2 |
| **Quercetin** | -9.9 | -5.0 | -10.6 | -10.7 | -6.4 | -9.3 | -11.6 |
| **Terminic acid** | -12.6 | -7.5 | -13.7 | -16.2 | -9.0 | -14.6 | -13.1 |
| **Catechin** | -8.6 | -3.9 | -8.9 | -9.1 | -5.6 | -5.5 | -10.0 |
| **Rutin** | -7.2 | -4.7 | -8.3 | -9.6 | -7.0 | -6.0 | -7.7 |
| **Kaempferol** | -8.5 | -4.6 | -8.5 | -9.3 | -5.6 | -5.9 | -7.5 |
| **Leucocyanidin** | -8.8 | -5.0 | -7.9 | -8.8 | -5.8 | -5.5 | -7.1 |
| **3-O-Methylelagic acid 3'-rhamnoside** | -7.5 | -5.0 | -9.0 | -10.3 | -6.1 | -5.9 | -7.4 |
| **Oleanolic acid** | -6.6 | -4.4 | -7.1 | -16.2 | -6.2 | -6.2 | -7.2 |
| **Inhibitors** | -4.2<br>(Phosphocholine) | -7.2<br>(Simvastatin) | -9.4<br>(Rofecoxib) | -9.6<br>(Carazolol) | -6.0<br>(Lisinopril) | -9.7<br>(Losartan) | -11.1<br>(Losmapimod) |

It was found that all the phytocompounds showed strong binding with receptor 1B09 which is higher than that of drug PC (−4.2kcal/mol). Arjungenin (−16.1 kcal/mol), casuarinin (−15.4 kcal/mol), arjunic acid (−13.3 kcal/mol), terminic acid (−12.6 kcal/mol) showed higher binding energy with 1B09. With 1HW9 receptor, casuarinin (−12.0 kcal/mol), arjungenin (−8.4 kcal/mol), terminic acid (−7.5 kcal/mol), arjunic acid (−7.4 kcal/mol) showed higher binding energy as compared to drug simvastatin (−7.2 kcal/mol). Casuarinin (−19.4 kcal/mol), arjungenin (−17.3 kcal/mol), arjunic acid (−14.3 kcal/mol), terminic acid (−13.7 kcal/mol), showed higher binding energy with 1cx2 receptor, as compared to drug rofecoxib (−9.4 kcal/mol). Casuarinin (−24.0 kcal/mol), arjungenin (−19.3 kcal/mol), arjunic acid (−16.9 kcal/mol), arjunolic acid (−16.8 kcal/mol), showed higher binding energy with 4YCW receptor, as compared to drug Carazolol (−9.6 kcal/mol). Casuarinin (−14.7 kcal/mol), terminic acid (−9.0 kcal/mol), arjungenin (−8.4 kcal/mol), arjunic acid (−7.1 kcal/mol) showed higher binding energy with 1O8a receptor, as compared to drug lisinopril (−6.0 kcal/mol). Casuarinin (−21.0 kcal/mol), arjungenin (−16.5 kcal/mol), terminic acid (−14.6 kcal/mol), arjunetin (−10.3 kcal/mol) showed higher binding energy with 4YAY receptor, as compared to drug losartan (−9.7 kcal/mol).With 4dli receptor, arjungenin (−21.6 kcal/mol), casuarinin (−17.0 kcal/mol), terminic acid (−13.1 kcal/mol), ellagic acid (−11.8 kcal/mol) showed higher binding energy with 1B09 as compared to Losmapimod (−11.1 kcal/mol) (Table 3).

### Drug-likeness prediction

The Drug-likeness filters help in the early preclinical development by avoiding costly late step preclinical and clinical failure. The drug-likeness properties of molecules were analyzed based on Lipinski rule of 5(Table 4). All the selected phytocompounds satisfied Lipinski’s rule of five, except casuarinin.

**Table 4:**
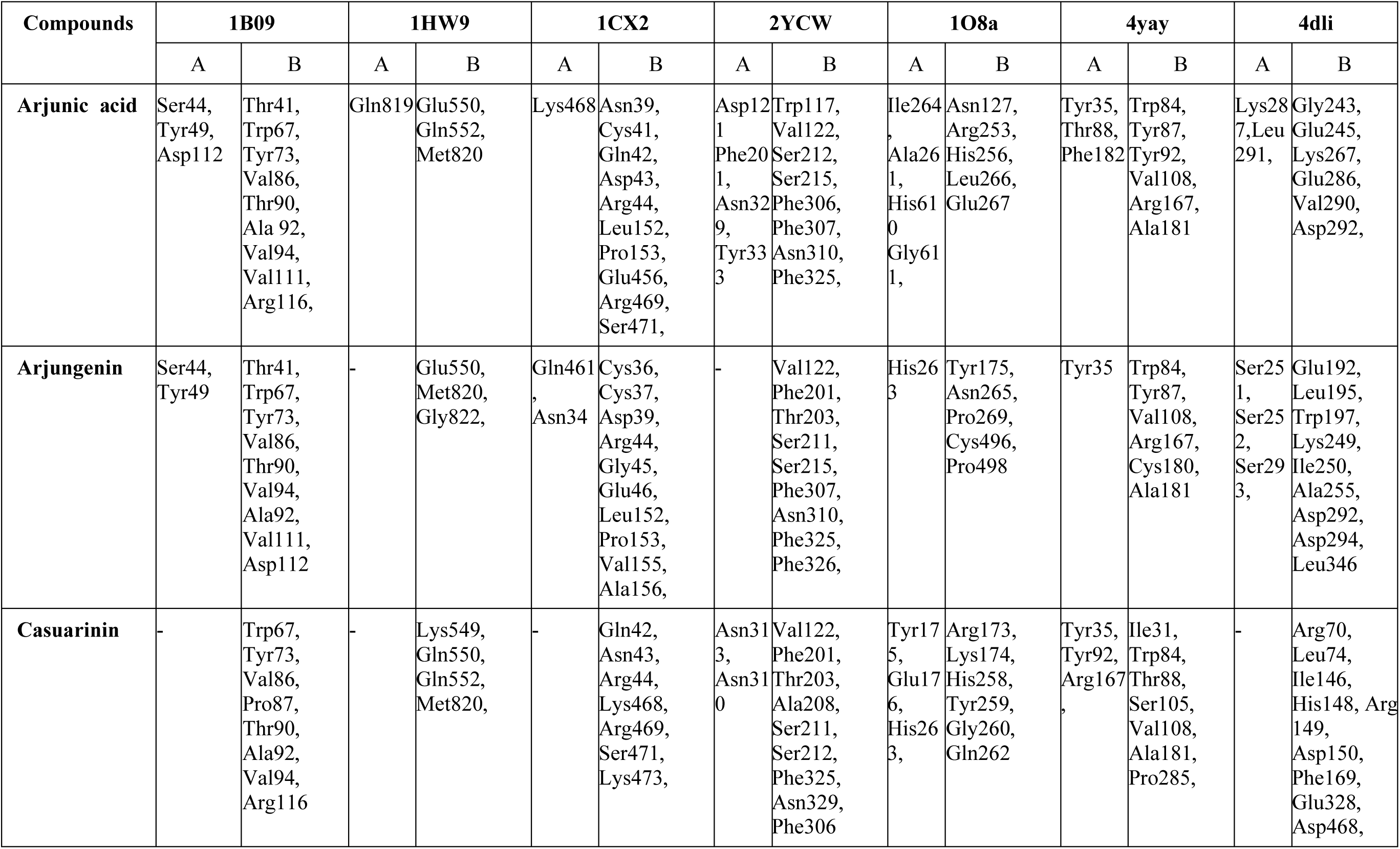
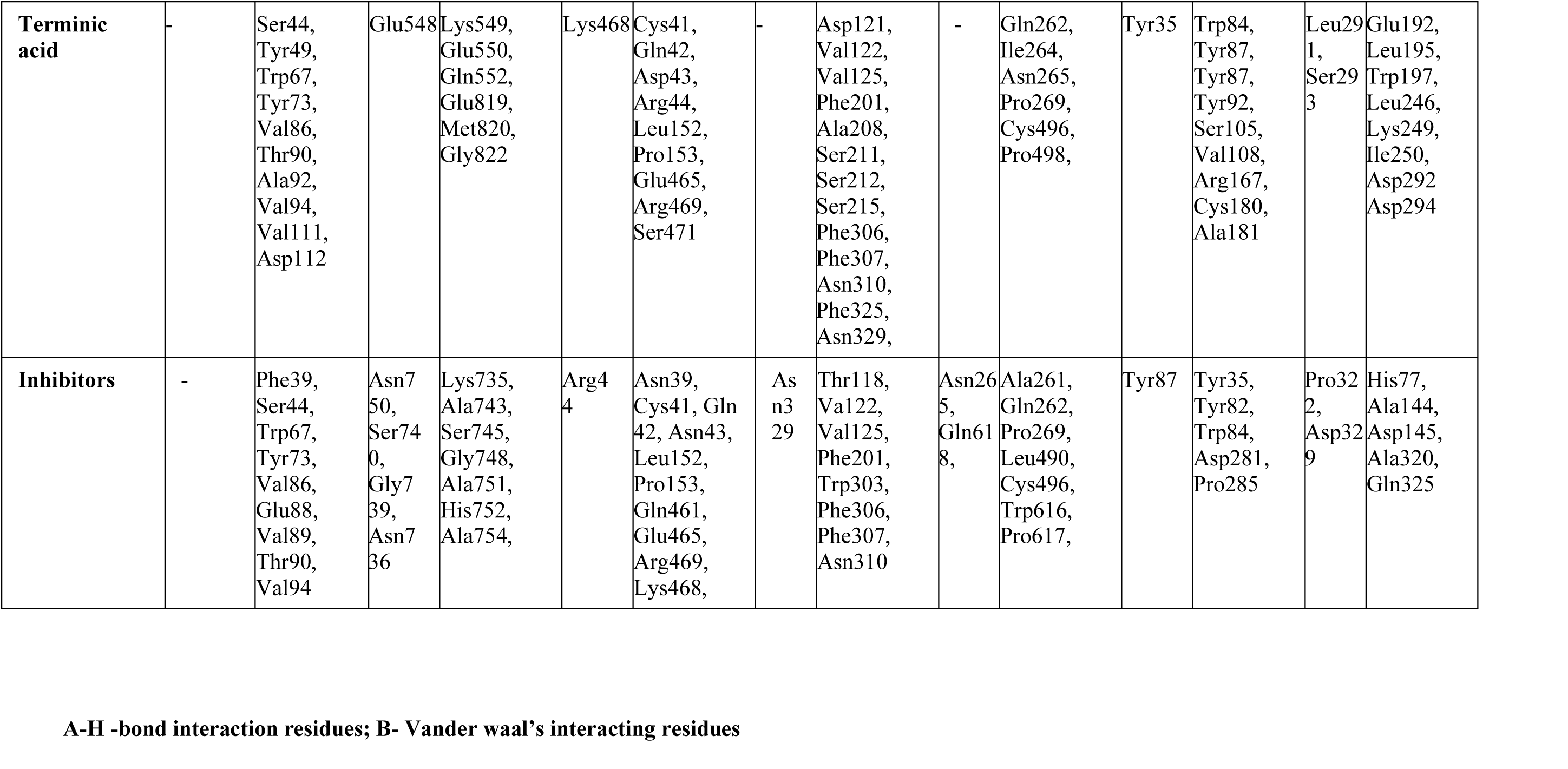
Intermolecular H-bonding and hydrophobic interaction between phytocompounds (Arjunic acid, arjungenin, casuarinin, terminic acid, standard drugs (inhibitor) and target proteins.

**Table 4:**
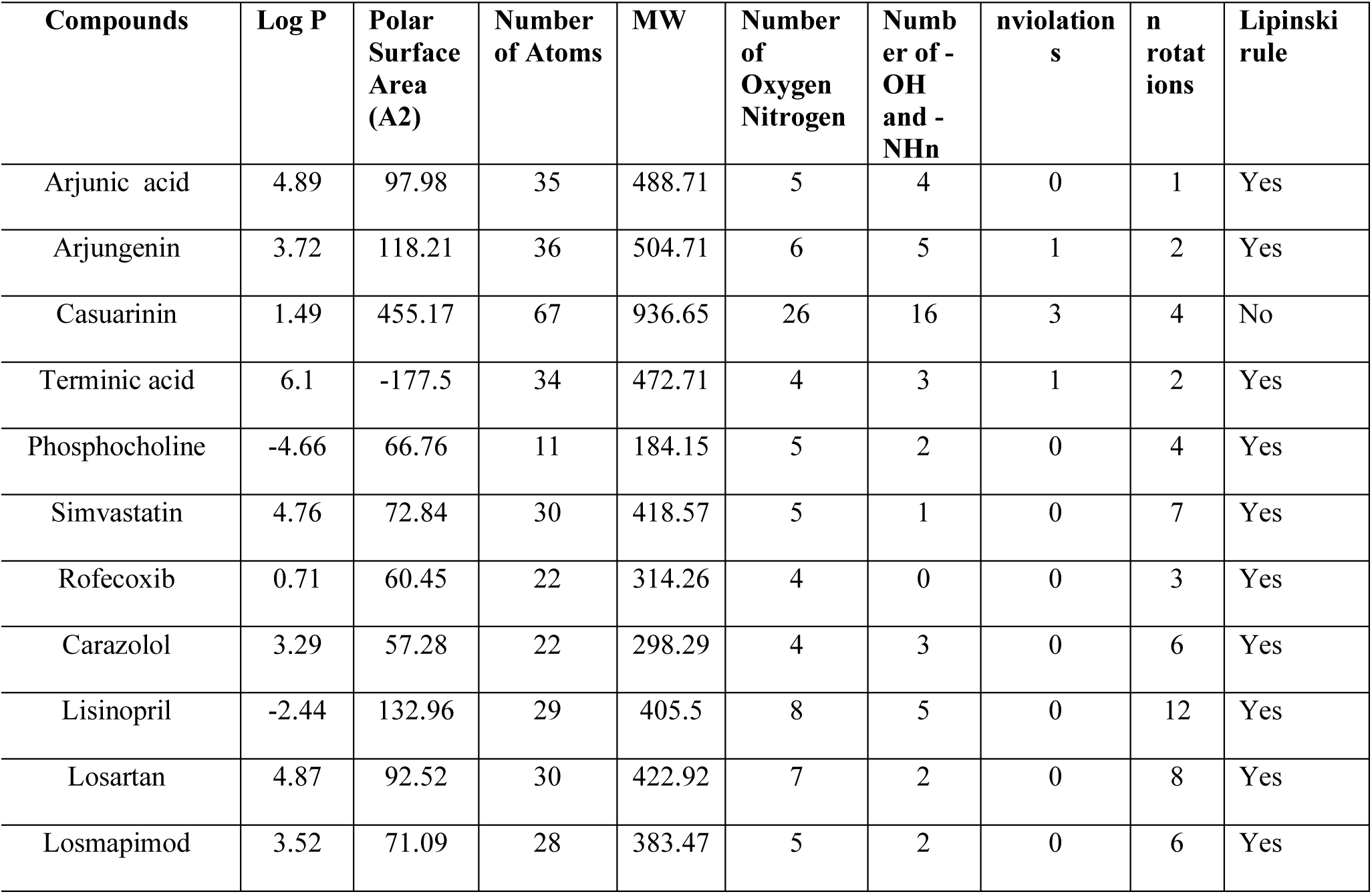
Drug-likeness prediction of selected phytocompounds from *T. arjuna*.

### Toxicity and ADMET prediction

Toxicity of phytocompounds were analyzed through Protox II server. The server admetSAR generates pharmacokinetic properties of compounds under different criteria: Absorption, Distribution, Metabolism, and Excretion [32]. The results of admetSAR analysis and toxicity prediction have been shown in table 5.

**Table 5:**
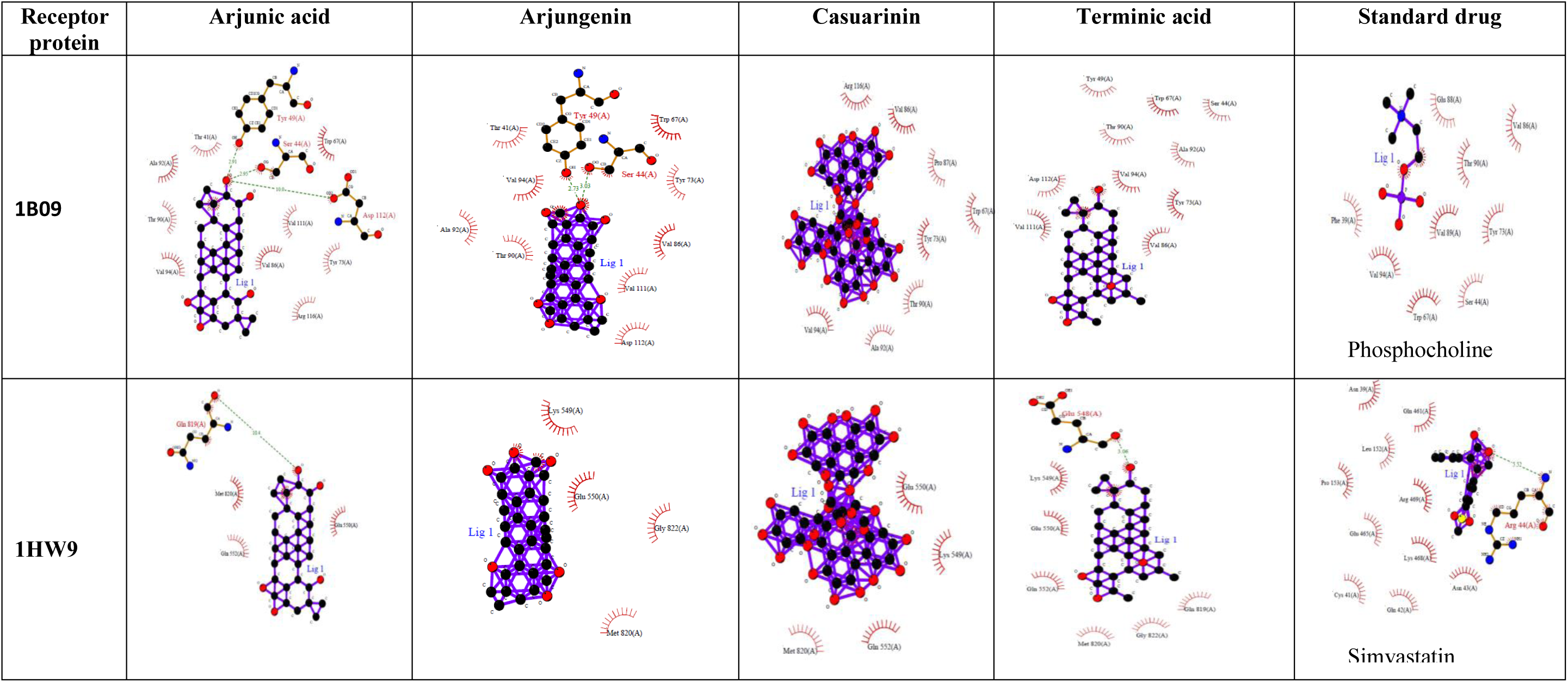
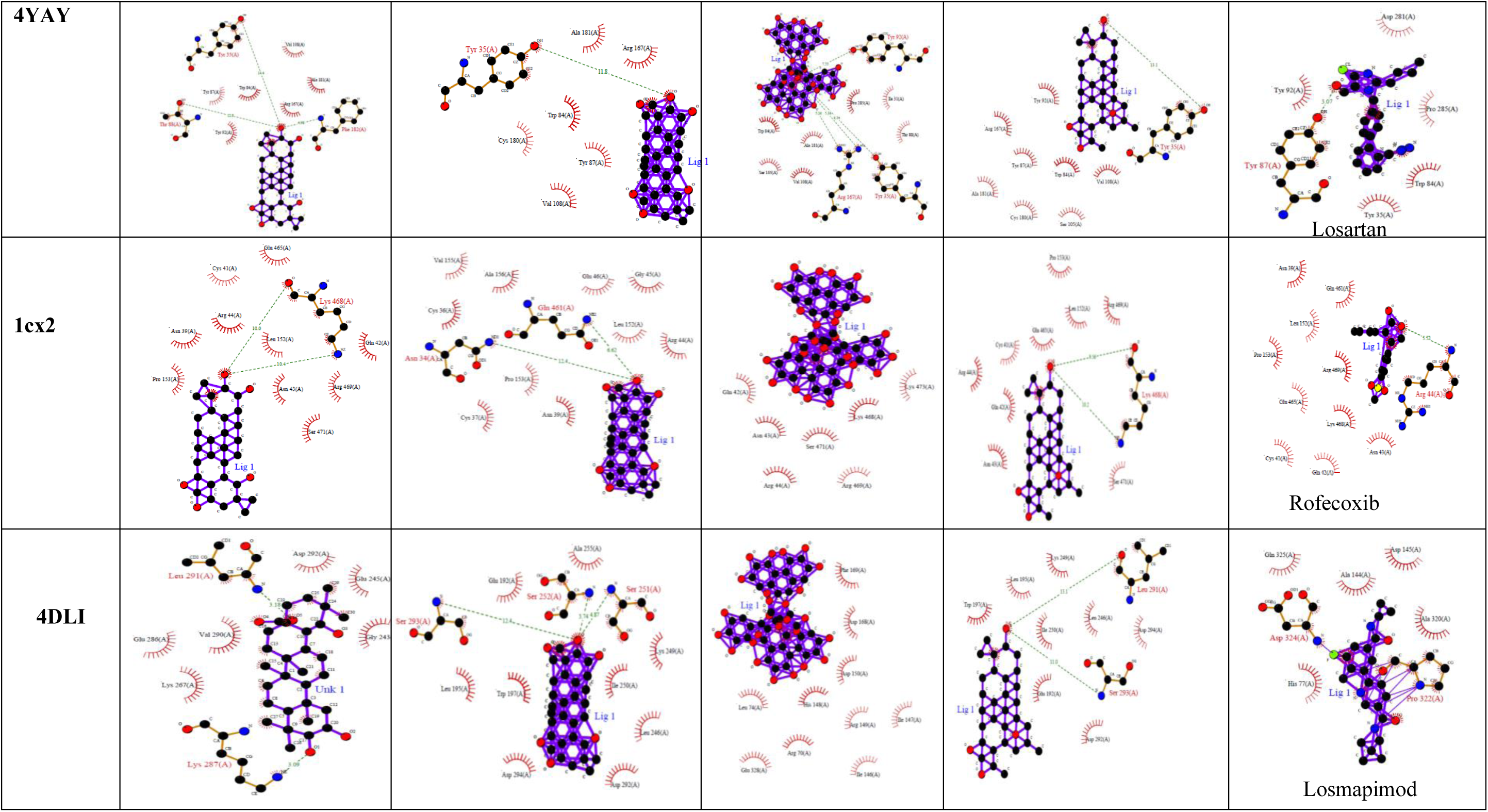
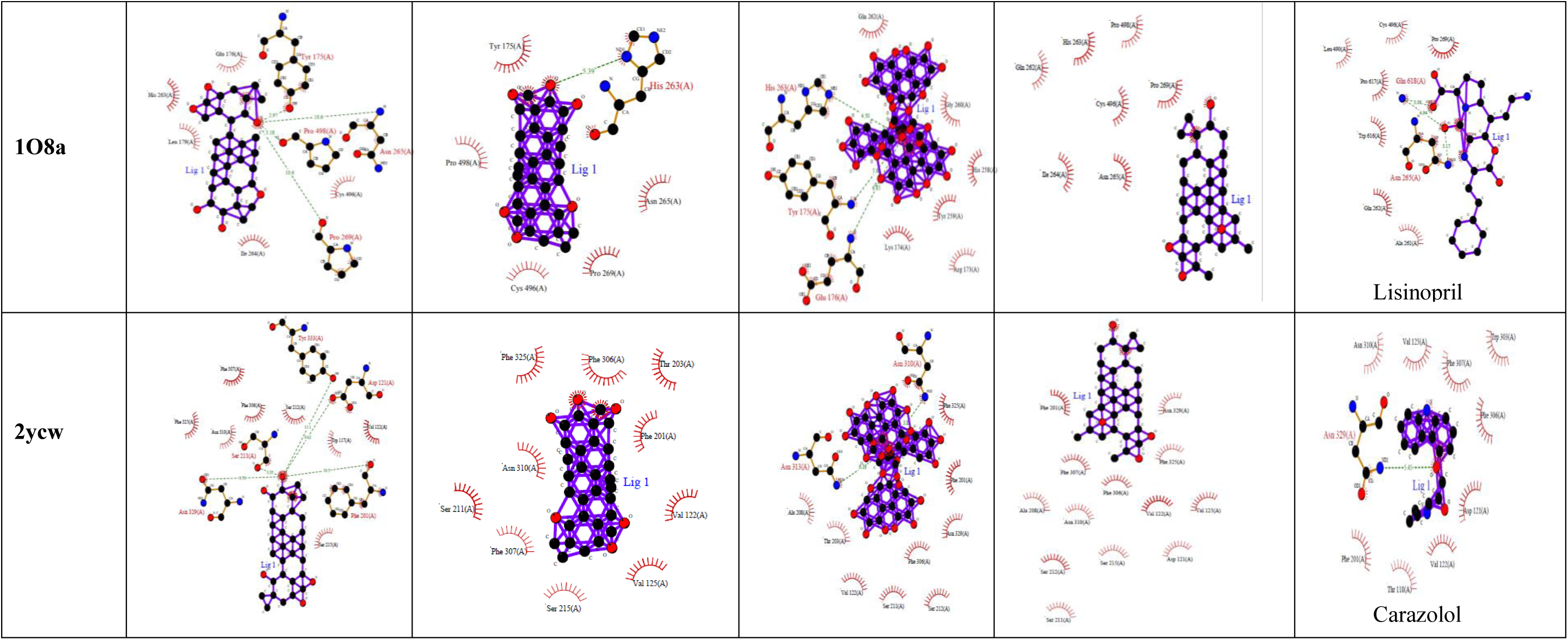
LigPlot structure showing interactions between phytocompounds, drugs and target proteins.

**Table 5:** ADMET and Protox-II prediction of selected phytocompounds of *T. arjuna* and drugs used through Admet SAR and Protox-II software.

| Compounds | Admet SAR |  |  | Protox-II |  |  |
| --- | --- | --- | --- | --- | --- | --- |
|  | Human intestinal absorption | Carcinogens | Rate Acute toxicity (LD50) kg/mol | LD50, (mg/kg) | Hepatotoxicity | Cytotoxicity |
| Arjunic acid | Positive | non-carcinogen | 2.71 (III) | 2000 (Class 4) | Inactive | Inactive |
| Arjungenin | Positive | non-carcinogen | 2.806 (III) | 2000 (Class 4) | Inactive | Inactive |
| Casuarinin | Positive | non-carcinogen | 1.828 (I) | 1000 (Class 4) | Inactive | Inactive |
| Terminic acid | Positive | non-carcinogen | 4.629 (III) | 2500 (Class 5) | Inactive | Inactive |
| Simvastatin | Positive | non-carcinogen | 2.301 Class III) | 1000 (Class 4) | Inactive | Inactive |
| Rofecoxib | Positive | non-carcinogen | 2.653 (Class III) | 4500 (Class 5) | Inactive | Inactive |
| Carazolol | Positive | non-carcinogen | 3.179 (Class II) | 235 (Class 3) | Inactive | Inactive |
| Lisinopril | Negative | non-carcinogen | 2.438 (Class III) | 8500 (Class 6) | Inactive | Inactive |
| Losartan | Positive | non-carcinogen | 3.322 (Class III) | 300 (Class 3) | Inactive | Inactive |
| Losmapimod | Positive | non-carcinogen | 1.611 (Class III) | 200 (Class 3) | Inactive | Inactive |
| Phosphocholine | Positive | non-carcinogen | 2.807 (Class III) | 1640 (Class 4) | Inactive | Inactive |

All of the phytochemicals showed an acceptable range of ADMET profiles that reflect their efficiency as potent drug candidates. All the compounds showed good human intestinal solubility (HIA), and acute rat toxicity (LD_50_) of selected compounds also belongs to same class (Class-II and Class III) to that of drugs except casuarinin (Class-I). None of the compounds are carcinogenic (Table-5). All the selected phytocompounds are inactive for cytotoxicity and hepatic toxicity. LD50 value for all selected compounds was higher except casuarinin (1000 mg/kg), indicating non toxic nature of these compounds. Therefore, based on ADMET and toxicity analysis, phytochemicals such as arjunic acid, arjungenin, terminic acid fulfill all the enlisted criteria and we can suggest that these are potential candidates for the development of a better drug against cardiovascular diseases.

## Discussions

In pharmaceutical research, computational strategies are of great value as they help in the identification and development of novel promising compounds especially by molecular docking methods [33, 34]. Various research groups have applied these methods to screen potential novel compounds against a variety of diseases [35]. Angiotensin-converting enzyme (ACE) has a significant role in the regulation of blood pressure and ACE inhibition with inhibitory peptides is considered as a major target to prevent hypertension [35]. Several studies have used docking approach to inhibit the expression of ACE protein with natura compounds. Quercetin glycosides showed optimum binding affinity with angiotensin-converting-enzyme (−8.5 kcal/mol) as compared to enalapril (−7.0 kcal/mol), thereby indicating that Quercetin glycosides could be used as potential candidate to treat hypertension, myocardial infarction, and congestive heart failure [36]. Similarly, methyl gallate and quercetin 3-*O*-β-D-glucopyranosyl-(1’’’-6’’)-α-rhamnoside are potential active compounds as ACE inhibitors from *P. niruri* herb [37]. Khan [38] studied the interactions between 4YAY (Angiotensin-I) receptor protein with phytocompounds from *Alangium salvifolium*. The compound alangum1 [Alangium1(4(benzoyloxy) methyl-2hydroxyphenoxy tetrahydorxy hexoxone 1,2,3,4,5, pentaium] showed the best glide docking XP score −8.5 kcal/mol binding energy value with best fit simulation study. Impertonin from whole fruit extract of *Aegle marmelos* showed strong interaction with HMG-CoA reductase enzyme [39]. Secondary metabolites such as Dichloroacetic acid 2, 2-dimethylpropyl ester, 1, 6, 10-Dodecatriene-3-ol, 3, 7, 11-trimethyl-[S-(Z)]-, Isopropyl acrylate and 3, 3-Dimethylacryloyl chloride formed strong binding with active sites of HMGCoA reductase [40]. Study from Sellapan *et al*. [41] reported better docking of plant derived compound, guajavarin with HMG-CoA protein. Study on phytocompounds from *T. arjuna* with phosphodiesterase 5A, sodium-potassium pump and beta-adrenergic receptor showed that casuarinin showed multiple inhibitions on phosphodiesterase 5A and sodium-potassium pump, whereas Pelargonidin on phosphodiesterase 5A and beta-adrenergic protein targets [42]. Talapatra *et al*. [43] reported that phytocompounds from *Calotropis procera* such as methyl myrisate (−3.0) and methyl behenate (−3.2), β-sitosterol (−5.6), uzarigenin (−5.5) and anthocyanins (−5.4) showed good binding with CRP receptors. Caffeic acid had remarkable interaction with proteins involved in inflammatory response (COX-2, COX-1, FXa and integrin αIIbβIII), thereby, having the potential to be developed as cardiovascular-safe anti-inflammatory medicine [44]. In the present study, we have selected 19 compounds reported from stem bark of *T. arjuna* against protein targets of various cardiovascular disease. It was found that among all selected phytocompounds, arjunic acid, arjungenin, terminic acid satisfy all parameters of ADMET and toxicity, also showed good affinity with selected protein targets, therefore, they could be used as potential broad -spectrum candidate for treatment of different heart problems.

## Conclusions

The present work was an attempt to computationally identify phytocompounds from T. arjuna which can bind to the various targets of cardiovascular disease. The docking scores, analysis of the interactions of the compounds suggest that most of the compounds have the ability to bind to multiple targets involved in cardiovascular disease. ADMET and toxicity prediction showed arjunic acid, arjungenin, terminic acid could be used as potential candidate against cardiovascular disease.

## Acknowledgement

The authors acknowledge Shoolini University, Solan, for providing infrastructure support to conduct the research work.

## Declaration of competing interest

The authors declare that there are no conflicts of interest.

## Fundings

This research did not receive any specific grant from funding agencies in the public, commercial, or not-for-profit sectors.

## Notes

### Competing Interest Statement

The authors have declared no competing interest.

